# Synergizing Geometric Deep Learning and Data-Centric Methods for Improved Protein Structural Alignment

**DOI:** 10.1101/2023.07.14.549112

**Authors:** Daniel T. Rademaker, Jarek M. van Dijk, Farzaneh M. Parizi, Li C. Xue

**Affiliations:** Medical BioSciences, Radboud University Medical Center, 6525 GA Nijmegen, The Netherlands

**Author notes:** These authors contributed equally (Daniel T. Rademaker, Jarek M. van Dijk).

## Abstract

**Motivation:** Structures are replacing the role of sequences. Traditional bioinformatics research focused on sequences because they were easily obtained. Advances in techniques like cryo-electron microscopy, molecular modeling, docking algorithms, and structure prediction software have shifted the focus to structures. Given the importance of deep learning in many of these breakthroughs, it makes sense to also explore how it can modernize classic bioinformatics tools. However, empirical findings have shown that machine learning based methods have many pitfalls resulting in overoptimistic conclusions, including data leakage between test and training data. Thus, there is a need for new innovations to make neural networks more intelligible.

**Results:** We have developed vanGOGH, a geometric deep learning-based structural alignment approach that performs on par with the state-of-the-art without ever having been trained on a pair of naturally found homologs. We adopted a data-centric approach to address deep learning and data limitations by augmenting protein templates into synthetic homologs for training.

Our method allowed us to supplement homolog data by knowledge-driven augmentation, self-learning of relevant structural features by supervised examples and protein alignment that is competitive with state-of-the art methods.

**Availability:** GNN framework: https://github.com/DeepRank/deeprank-core/tree/main/deeprankcore

## Introduction

### Structural alignment

Protein alignment is a classic bioinformatics problem that has proven to be invaluable for a wide range of research areas. Its goal is to determine which amino acids in two or more homologous proteins are equivalent with respect to protein structure and function. Homologs are a result of mutation events after various possible differentiation mechanisms, such as gene duplication and speciation (orthologs), gene duplication within a species (paralogs), or convergent evolution (analogous). Numerous differences between homologs may occur, such as insertions, deletions a and substitutions of secondary structure elements (SSEs) as well as circular permutations, strand invasions/withdrawals and hairpin flips/swaps[1]. Since homologous proteins typically share a similar sequence, structure, and function, homology can be used to infer evolutionary relationships between different species and aid in identifying protein function from its homologs. Given that protein structure is more conserved than sequence during evolution, the structural information component is crucial for obtaining reliable alignments between more distantly related proteins. However, structural alignment is a NP-hard problem, which means that an optimal solution cannot be guaranteed and the solution will always rely on heuristics[2].

There exists a plethora of structure alignment methods, each with its own set of advantages and disadvantages. The PyMOL [3] molecular visualization software provides an explicit example of tradeoffs in alignment methods, as the documentation of the ‘Cealign’[4] method states that it is robust for proteins with little to no sequence similarity, while the ‘super’ command is preferred for proteins with decent structural similarity, and the ‘align’ command is preferred for proteins with decent sequence similarity. However, tradeoffs can also be purely structural. Most methods have a performance bias to α-α and α-β proteins (CATH classification), while the β-β class is underrepresented[5]. In contrast the LSQMAN[6] method overrepresents this latter class and underrepresents the α+β protein pairs[7]. DALI[8] has made a heuristic trade-off by being limited to globular proteins with a compact core of alpha-helices and/or beta-sheets, thereby limiting its use for small proteins[9]. Additionally, DALI defines similarity relative to intramolecular Cα distances thereby permitting larger deviations for tertiary contacts (between SSEs) than the local ones (such as backbone residue distances)[10]. Tradeoffs are also found in TMalign[9] in the use of its TM-score[10] during alignment. The score weights smaller distance errors more strongly than large ones, is sensitive to global similarities instead of local variation, and creates length-independency through normalization of distance errors[11-13]. Additionally, there is a difference in consistency of alignment between methods. FATCAT[14] for example, produces good alignment scores but shows inconsistency in its results, whereas SAP[15] is more consistent at the cost of accuracy[7]. In terms of consistency and performance, TM-align performs well for structurally similar domains, but is outperformed by Fr-TM-align for less similar domains[7].

Many folds will have their own patterns and rules and therein lies the problem. Although all these designed structural alignment methods are innovative and clever, a few handwritten rules can never adequately represent the diversity of patterns present in the many protein-folds. And rather than painstakingly extracting these patterns by hand, it may be more effective to employ a machine learning method to automatically extract them from the vast amount of available protein-structure data. Deep learning would be the ideal candidate for learning the patterns, and we believe the problem of structural alignment would be an ideal testcase to explore if the deep learning algorithms truly ‘understands’ the structural data.

### Deep learning

Deep learning is becoming a general-purpose algorithm whose results are starting to dominate many problem domains[13]. It’s part of a new programming paradigm where algorithms are learned rather than designed. The effectiveness of neural networks can be attributed to two factors: model architecture and training data. Although both important, most published deep learning research focuses mostly on improving neural architectures which are then tested on the same gold standard datasets.

The neural network architecture is often chosen based on the data modality, enabling engineers to incorporate domain knowledge about the problem directly into the neural network [13]. When done correctly, this reduces the search space, making the problem easier to solve for gradient descent optimization. Convolutional Neural Network (CNN) architectures, for example, are well adapted to deal with image data thanks to their ability to exploit translational symmetries. They have also been successfully applied to protein structural data. Their results, however, partially depend on arbitrary protein rotation in Euclidian space, which is an undesirable property of CNNs. As a workaround, researchers typically train CNNs on several rotations of the same input structure, thereby undesirably necessitating the need for more computing time and neural network parameters. A popular alternative is the use of Graph Neural Networks (GNNs) as they are naturally rotationally invariant while still remaining translationally equivariant, they can be a preferable alternative for managing protein data.

Thanks to efforts such as the ‘Geometric Deep Learning’ theory, which attempted to unify *the veritable zoo of neural network architectures*[13], the problem of neural architecture design has been essentially solved for most types of data. However, the impact of training data, which is on the other side of the proverbial deep learning coin, remains largely unexplored. Some researchers, most notably Andrew Ng, often speak of the “proof of concept to real-world production gap”[16] difficulty and appoint the training data as the limiting factor for neural network performance rather than the neural architecture design. This limitation often occurs in many real-world research fields, such as proteomics, where data can be noisy, small in size, unlabeled or mislabeled, and unevenly distributed. While researchers have attempted to overcome these limitations by tweaking the neural architecture and creating intricate training schemes, a growing alternative is to optimize the data itself. This approach, known as “data-centric learning,” is recently gaining traction as a means of improving performance on challenging datasets.

### Summarizing paragraph

In our research, we have combined synthetic data training and contrastive learning methods to investigate how data augmentation can facilitate the integration of domain knowledge into GNN networks, eliminating the need for explicit coding of that knowledge within the architecture. By leveraging a data-centric approach, our method minimizes the reliance on external labeled datasets, making it more adaptable and less susceptible to biases that may be present in such data. The outcomes of our research have yielded a highly competitive algorithm for structural alignment, which we have aptly named vanGOGH. This acronym stands for variational augmented neural Graph of Generalized Homology, encapsulating the essence of our approach.

## Materials and methods

### Data download and preprocessing

To obtain the dataset for our study, we downloaded 33,694 protein domains from the Structural Classification of Proteins (SCOP) database [17], [18]. Next, we used the DSSP tool [19], [20] to segment each domain into simplified secondary structural elements (SSEs), which were classified as helices, strands, or loops (see Figure 1). Since the backbone is more rigid than the sidechains, here, and in nearly all structural alignment algorithms, each amino-acid was reduced to the Cα atom from the protein backbone. For more detailed information on this process, we refer to the supplementary materials.

**Figure 1.**
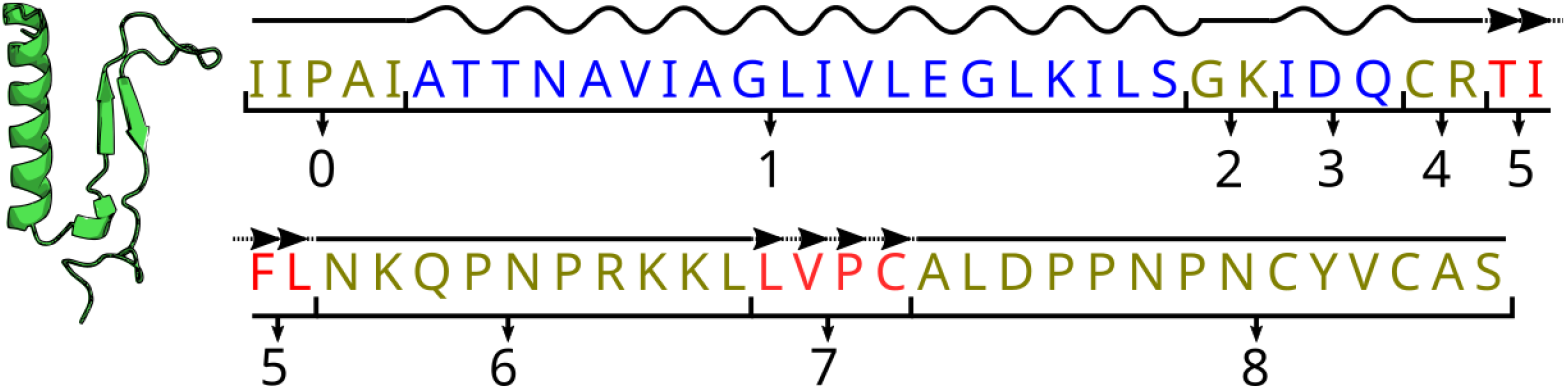
Illustration of the transition of protein to node and edge features. Each amino acid is assigned one of three secondary structural elements (helix in blue; sheet in red; loop in yellow) using the DSSP method and each SSE is numbered.

### Neural architecture and features

The GNN architecture used in this study is defined by equations 1 through 6, of which the notation style was inspired by Satorras *et al*.[21]. Non-linear operations (∅) are represented by Multilayer Perceptrons (MLPs), with each ∅ having a single hidden layer, except for ∅_h_, which has two hidden layers due to the residual connection in equation 5. In this study, each MLP had a hidden layer with 32 neurons.

GNNs receive input in the form of node and edge features. For our GNN the node features secondary-structure type (**u**) and amino-acid type (**y**) per residue were represented as one-hot vectors. The edge features between neighboring nodes *i* and *j* contain four types of information: the squared distance (**d**) in angstroms, whether or not nodes *i* and *j* belong to the same secondary-structure element/stretch (**s**) (Figure 1), if nodes *i* and *j* are direct neighbors in the protein-sequence (**c**), and if node *i* and node *j* in the sequence encodes information on the relative position of the two nodes, specifying whether node *i* is preceding node *j* or vice versa in the sequence (**r**). Both node and edge features are transformed into node (**h**) and edge (**a**) embeddings (eq. 1-2) before entering the GNN layers, with an embedding size of 32 for each.

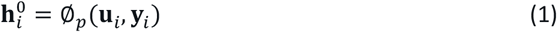

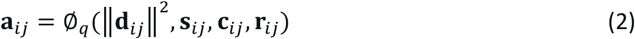

GNN layers are defined using equations 3-6. Messages (**m**_*ij*_) are calculated between neighboring for each pair of nodes with a distance less than 15 angstrom (eq.3), and subsequently summed per node (eq. 4). The resulting **m**_*i*_ are then used to update the node embeddings (eq. 5).

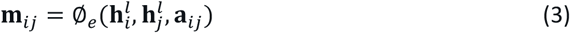

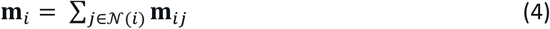

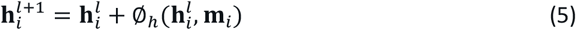

The final GNN layer is used to generate the final output embedding (**o**_*i*_) per residue, which encodes the processed local structural and amino-acid information. In our study this embedding had a size of 32.

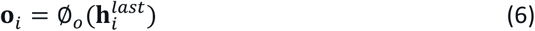

The local context size on which the output vectors are produced is dictated directly by the radius used to connect the nodes and indirectly by the number of layers. Given enough layers, the neural network can learn itself how much local context to utilize for creating its representations. However, it is important to keep in mind that the neural capacity may also have an impact on how much information the GNN can absorb. To ensure that we can solely focus on the input data, we provided the GNN architecture with ample capacity and layers so that we did not need to adjust the architecture itself.

Through the development of the GNN’s features, we have adjusted the features multiple times to improve performance. To show the significance of feature engineering, we also implemented our approach with simpler version of the GNN that omits the “same strand” (**s**) and “relative positions” (**r**) features. In the results section this model will be referred to as “GNN_basic”, whereas the GNN incorporating all features is referred to as “GNN_full.” When the GNNs are mentioned without specifying the features, it should be assumed that all features are utilized.

### Synthologs

To ensure that our neural network training had an equal representation of protein folds, we required a dataset of protein homolog pairs that were uniformly sampled from all 1,539 protein folds defined by SCOP. However, since many of these folds did not have naturally occurring homologs in the SCOP database, and obtaining new protein structures through experiments is quite impractical, we opted to augment existing protein structure templates to mimic natural homologs. In the remainder of this article, we refer to these synthetic homologs as “synthologs”.

To simulate the structural variations observed in homologs, we employed a stochastic selection and augmentation process for the simplified SSEs, as illustrated in Figure 2a. This involved introducing new orientations through a technique called “wiggling” of the segment. To determine the rotation axes for the wiggling, we utilized Principal Component Analysis (PCA) to identify the main axes of variation, as shown in Figure 2b. Rotations were then applied around PC1, PC2, and PC3, where PC1 corresponds to the rotation of an SSE around its central axis (such as the spiraling direction of a helix, as depicted in Figure 2b), while PC2 and PC3 represent shifts in the direction orthogonal to the central axis (Figure 2d).

**Figure 2.**
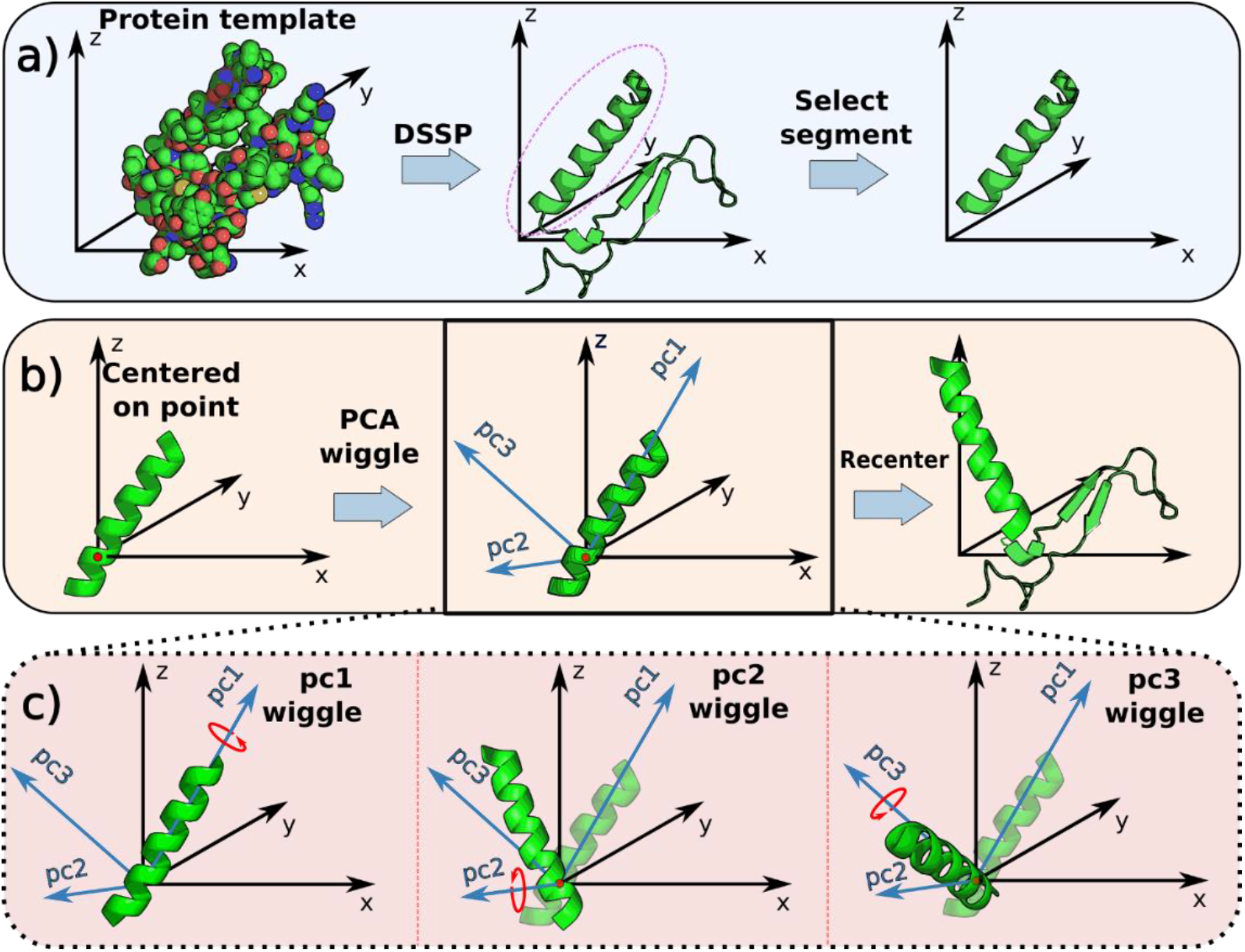
Procedure for augmenting protein domains. a) DSSP is used for SSE-based segmentation and the protein domain is reduced to the C-alpha backbone. b) A selection of SSE-segments are rotated using PCA estimates a segment’s internal orientation after random centering, followed by rotations over pc1, pc2 and pc3 and re-centering back to the initial location. C) PCA-based rotations

After performing the wiggling process on all SSEs, we introduced a small amount of noise to the Euclidean coordinates of the amino acids. This was done by adding random values between 0 and 0.1 angstrom to emulate the observation errors that often are present in low-resolution Protein Data Bank (PDB) structures. Given that we exclusively focused on the primary structure of proteins by considering backbone Cα residues, it was straightforward to simulate amino acid substitution. This involved replacing the amino acids in a syntholog according to its substitution score, which indicates the likelihood of evolutionary compatibility, as determined by the BLOSUM80 matrix. For more information on the specific process of replacement, please refer to the supplementary materials.

### Training the GNN

Our main assumption about the data is that corresponding residues between protein homolog pairs have similar structural and chemical local environments. During training the network was thus optimized to create similar embeddings for similar residue environments and dissimilar embeddings for dissimilar residue environments. Figure 3 shows the optimization procedure used to learn these embeddings. Initially two homologous proteins A and B are created from a template using structural augmentations of backbone proteins. To incorporate information on amino-acid similarity, the sequences of A and B in are substituted with likely amino acid alternatives (including no substitution) into A’ and B’. Additionally, protein B is copied and sequentially augmented to create a third protein C with an identical structure to B but with a very different sequence (unlikely mutations) compared to A’ and B’. To ensure similarity between embeddings of equivalent residues and dissimilarity between non-equivalent residues, we used the inner product between the embeddings of the homologous proteins as a distance measure. This inner product then results in a distance matrix between all residues between the two proteins (see Figure 3). We then applied a horizontal Softmax operation to this distance matrix, transforming each row into probability values representing the likelihood of each residue in one protein aligning with residues in the other protein. In the ideal scenario, the distance matrix between A’ and B’ would match the identity matrix of the same shape, which is why we used it as our optimization objective. This approach guarantees that equivalent residue embeddings are encouraged to be similar (towards the 1 labels), disregarding differences observed among homologous positions, while remaining dissimilar to other residues (towards the 0 labels).

**Figure 3.**
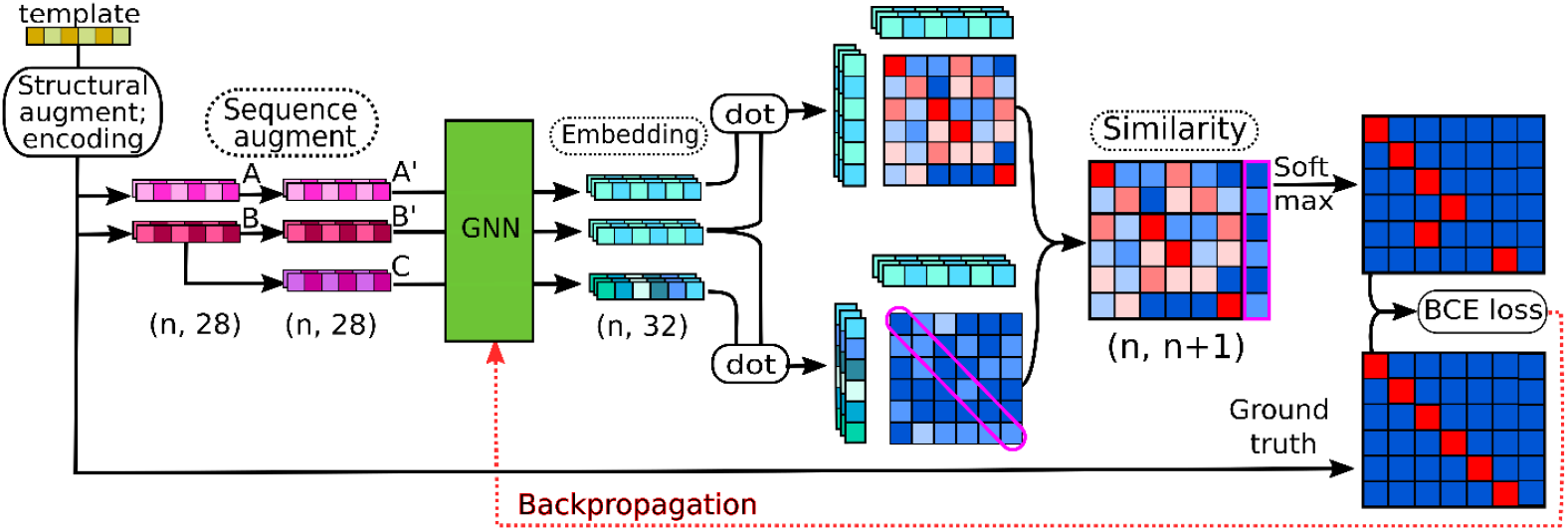
Training scheme of the GNN. A template PDB (yellow-green) is structurally augmented into two new structures (A/B), both encoded by One-Hot vectors for amino acids and node features (magenta/pink). The preliminary synthologs A and B are sequentially augmented into A’and B’ using probable augmentations, while B is also modified with improbable mutations into syntholog C. By concatenating the dot-product based similarity of embeddings (cyan) B’/C to the similarity of A’/B’, followed by a SoftMax function on the resulting matrix, we forced similarity between A’/B’ by a Binary Cross Entropy loss relative to the ground truth alignment (i.e. the identity matrix with an empty column attached).

However, this approach would not effectively learn the likelihood of different amino acid substitutions, as the contrastive learning approach is prone to making the GNN invariant to amino acid types. To address this limitation and ensure the consideration of both structural and chemical information, we introduced a second distance matrix between B’ and the structure C, which contains highly unlikely amino acid substitutions compared to B. This matrix specifically captures differences in residue type (chemical information) while keeping the structural information constant. To incorporate this information into the training process, we concatenated the diagonal of this second distance matrix with the original distance matrix, resulting in a final matrix with a shape of (n, n+1). By enforcing the embeddings of A to be dissimilar to those of C, the network is compelled to learn the distinguishing features between different amino acids. It learns which amino acids are similar and dissimilar, taking both structural and chemical environments into account. It’s important to note that this additional step is performed only during the training phase.

### Structural alignment

To align structures, it is necessary to calculate the correct rotation matrix that optimally superposes one protein onto the other (see Figure 4). This calculation is guided by the generated embeddings from the trained Graph Neural Network (GNN). Initially, the Kabsch algorithm [24] is employed to perform an initial raw structural alignment by pairing residues based solely on their highest similarity between the proteins. The alignments are then refined through 15 iterative steps. In each step, the median distance between the closest homologous residues is calculated. Subsequently, only the residues with a distance smaller than 1.5 times the median distance are aligned using the Kabsch algorithm once again. Through empirical observation, we have found these refinement steps to be necessary primarily for proteins that contain repeating motifs.

**Figure 4.**
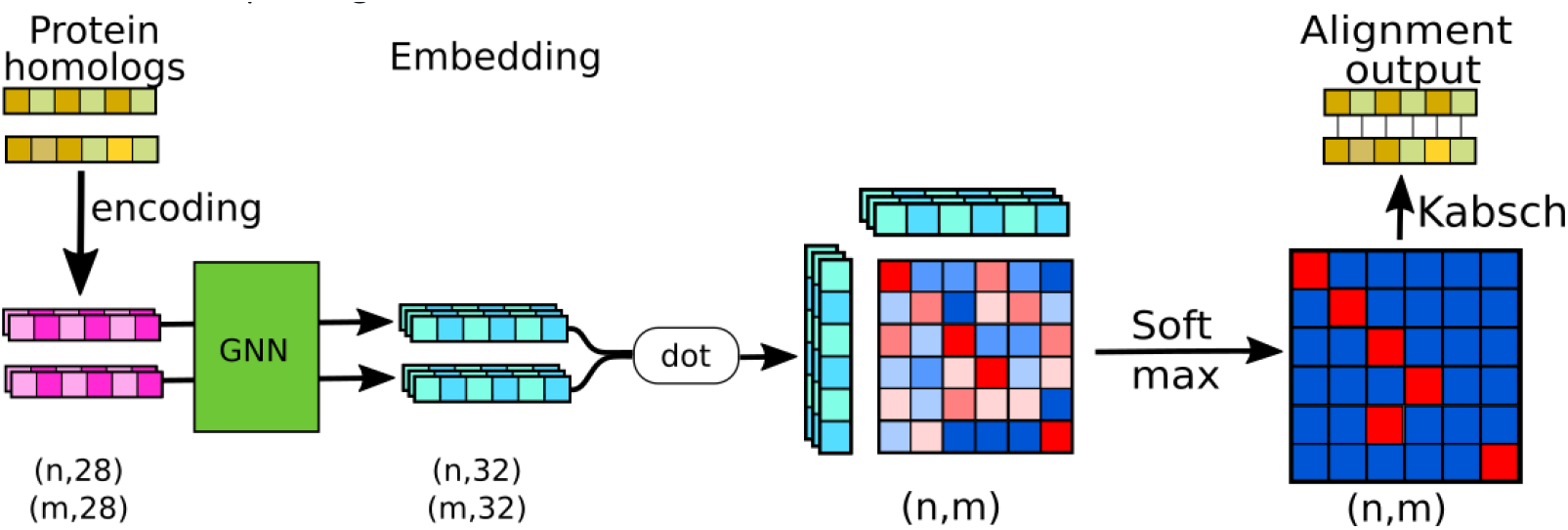
Alignment of protein structures. Similar to the training schema in Figure 3, the process begins with a GNN creating embeddings of feature-encoded proteins. Pairwise residue similarities are obtained through a dot/inner product operation on the embedded residues. These similarities are then translated into residue alignment probabilities using the Softmax function. Finally, the Kabsch algorithm is employed to optimize the alignment based on these probabilities.

## Results

### Scop distribution

Although there is a wide range of pdb data available (1539 folds spread over 33694 structures), there exists an uneven distribution of domains across the different folds, as depicted in Figure 5. Merely 63 folds (4.1% of the total) encompass a significant portion, accounting for 50% of the structures (16870 structures). Similarly, 185 folds (12%) cover 75% of the data, 444 folds (29%) encompass 90% of the data, and 677 folds (44%) represent 95% of the data. The remaining folds contain a median of approximately 4 structures per fold, with an average of 11.39 (std = 20.04, range = 1-126). Additionally, there were 419 folds (27.2%) that comprised only a single structure per domain, and two empty folds (which were subsequently removed from the dataset).

**Figure 5.**
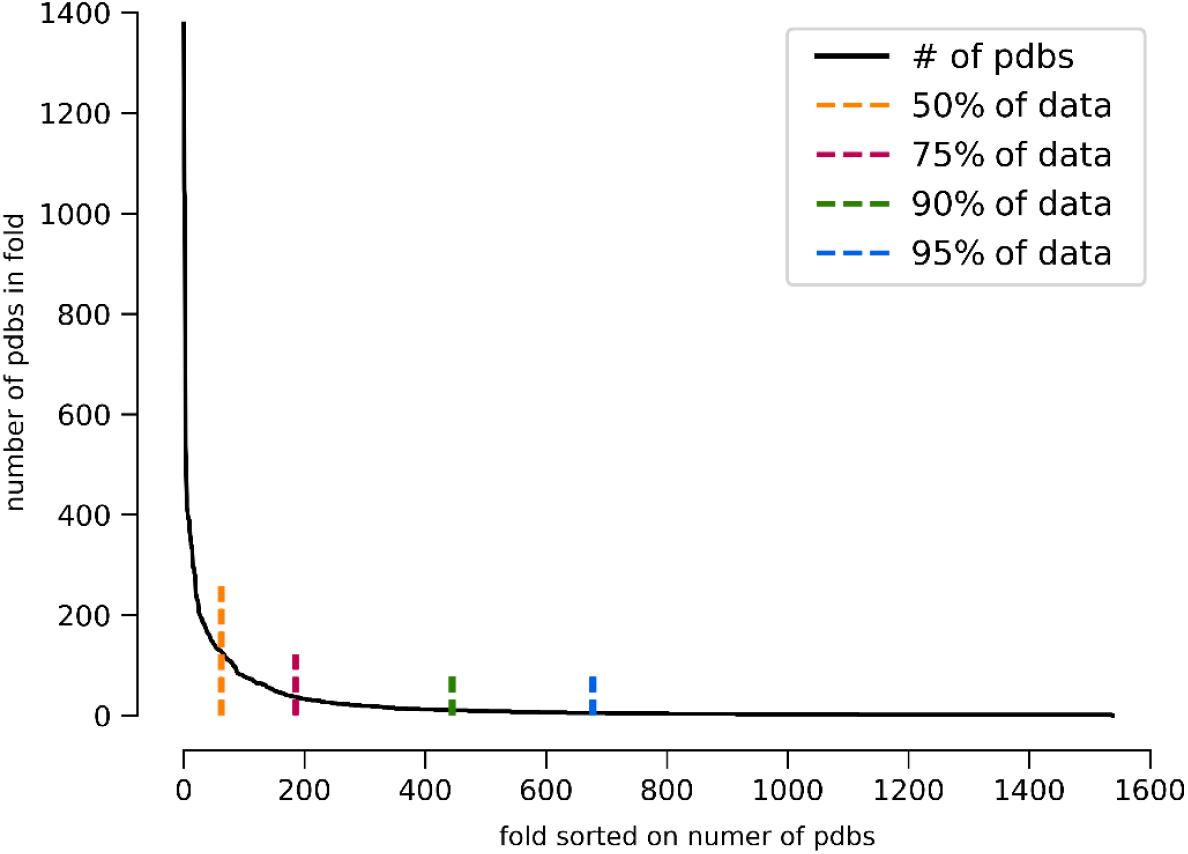
The non-uniform distribution of the proteins per SCOP class. The folds are sorted on the number of structures per fold. The black line shows the number of PDBs for the nth-fold (x-axis), whereas the broken lines indicate the percentage of data present before this fold.

### Synthologs

By employing the ‘wiggling’ technique, we generated synthologs that exhibit close structural resemblance, as illustrated in Figure 6. Despite originating from the same procedure and parameters, the observed differences among the wiggling outcomes demonstrate the capability of our approach to produce a diverse range of synthologs without requiring manual adjustments. Furthermore, by increasing the maximum allowed wiggling (as depicted in Figure 6b), we can enhance the variability of the synthologs, providing precise control over their sensitivity to variation.

**Figure 6.**
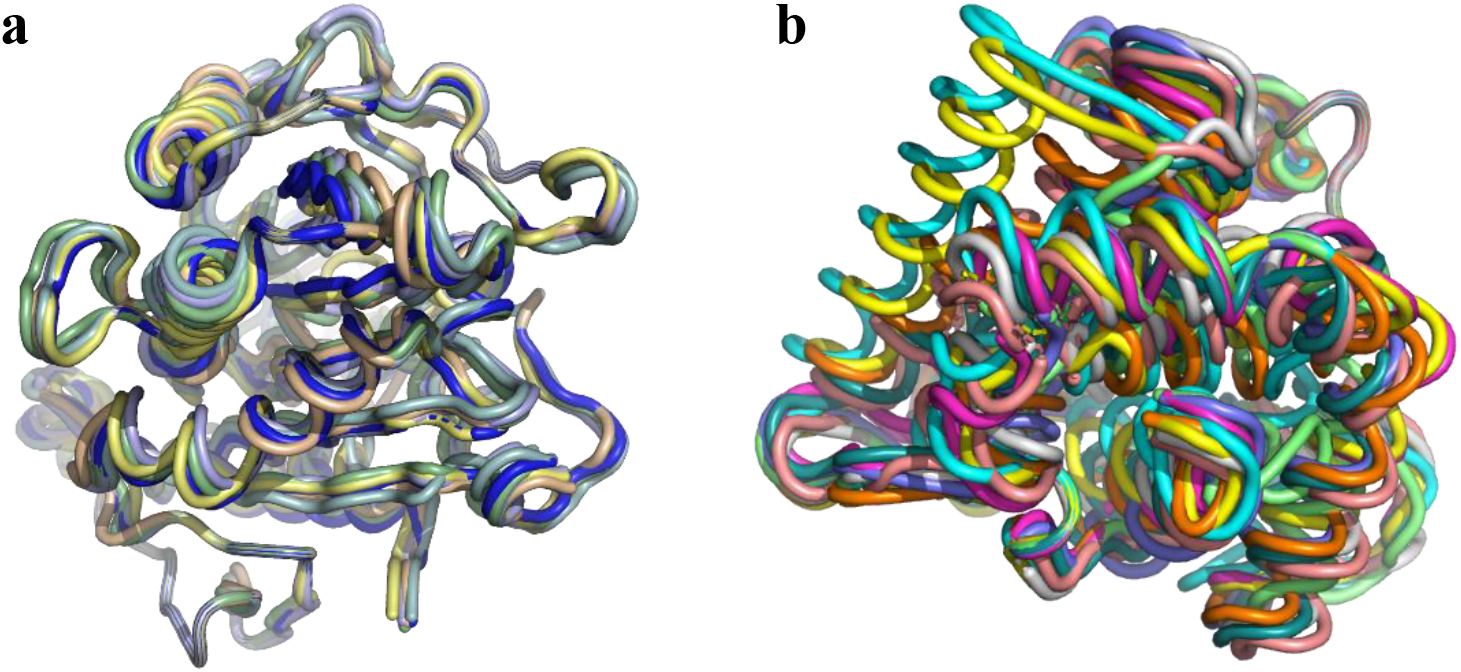
Several wiggling-based synthologs of 1GBS, using regular (**a**) and more extensive wiggling (**b**).

### Interpretability

To explore the learned GNN embeddings and assess whether they achieved our intended objectives, as well as to gain insights into what the model learned and identify the most important input features, we chose a visual approach. We employed an autoencoder to compress the 32-dimensional representation of the embeddings into three dimensions, bounded between 0 and 1 using a sigmoid activation function. This dimensionality reduction facilitated the visualization of the embeddings by adjusting the range to be compatible with various color spaces such as CIELAB and RGB. As a result, we were able to assign three color channels to each residue, enabling a visually interpretable representation of the embeddings. We refer to the supplementary materials for detailed information regarding the conversion process. Please note that the dimensionality reduction for coloring purposes may cause some loss of subtle differences between embeddings.

Initially, we analyzed the embeddings for *GNN_basic*, as comparing them to *GNN_full* provided insights into the impact of the additional features. During the analysis of the embeddings, we observed intriguing color patterns related to SSEs. Figure 7 demonstrates that the embeddings exhibit similar color patterns for helices (magenta-red), sheets (yellow-orange-green), and loops (blue-green). This consistent coloring suggests that the secondary structure input feature dominates the embeddings. Additionally, we found that the embeddings were resilient to small rotational variations between homologs as evidenced by the multiple variations of the same structure depicted Figure 7b indicating robust representations between equivalent residues.

**Figure 7.**
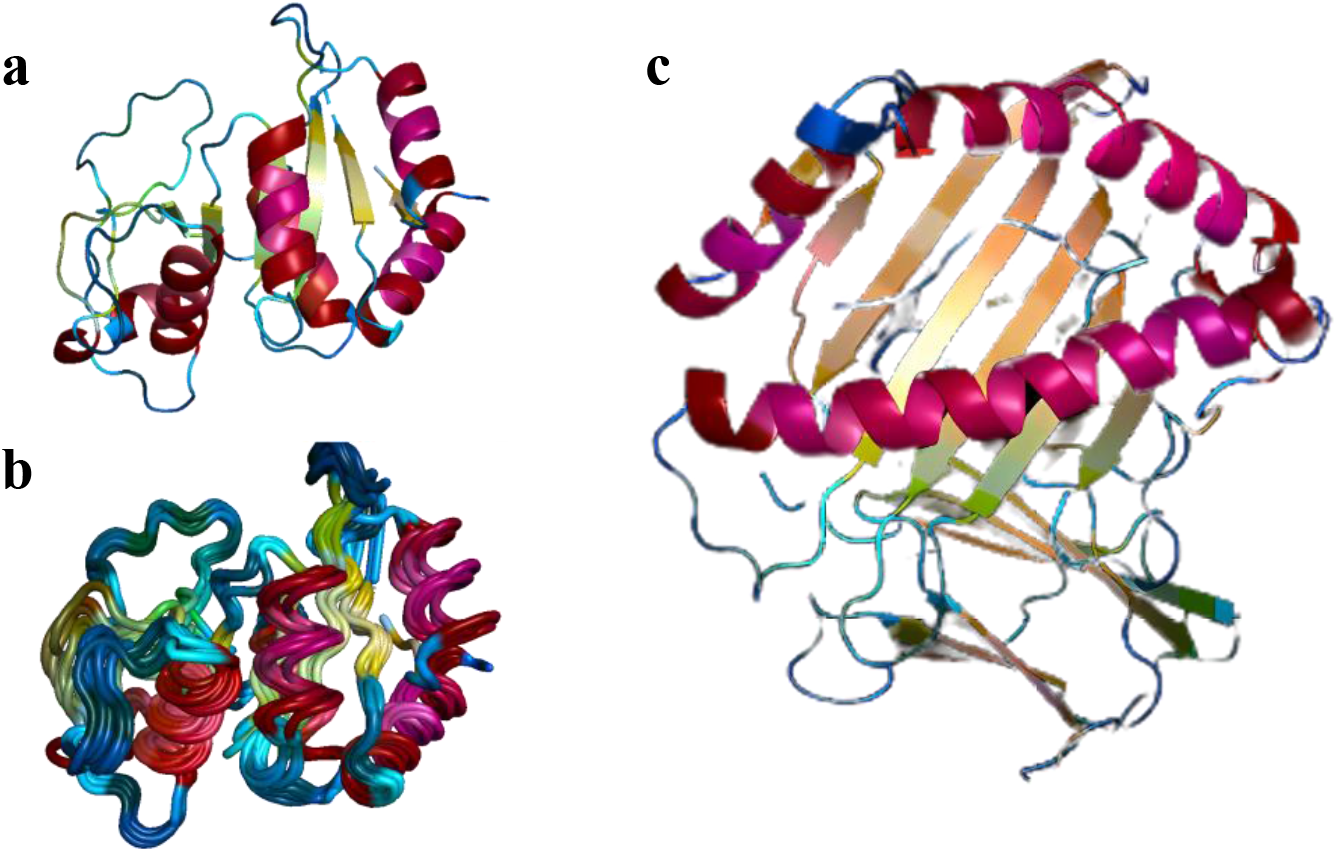
Visualization of learned GNN_basic embeddings. Dimensionality reduction is done with an autoencoder to three dimensions and translated into CIELAB colorspace. Panel (**a**) shows the embedding of 1G57. Panel (**b**) shows multiple synthologs created using this pdb as template. Panel (**c**) shows the embedding representation of MHC-I (1A1M).

While the embeddings displayed overall similarities between structures (Figure 7a and 7c) and same SSEs, we also observed subtle differences between them. For instance, in the MHC-I structure (Figure 7c), there are slight variations in color within the beta-sheet. The presence of orange color on the top right and green color on the bottom right of the beta-sheet suggests distinct chemical/structural environments within the same sheet.

When all features were incorporated during the training of the graph neural network (GNN_full), including information about sequence directionality and SSE identity, the embeddings changed accordingly. As shown in Figure 8, the embeddings became more homogeneous between different types of SSEs, indicating a better integration of information from non-secondary-structure features. However, each SSE still retained a distinct color gradient within the embeddings. For helices, the color gradient typically shifted from cyan to green and then to red. Similarly, loops displayed a gradient from cyan towards magenta, with more variation in the embedding space. The beta-sheet also exhibited a gradient from cyan to red, with a green-like color in-between for each individual strand of the sheet.

**Figure 8.**
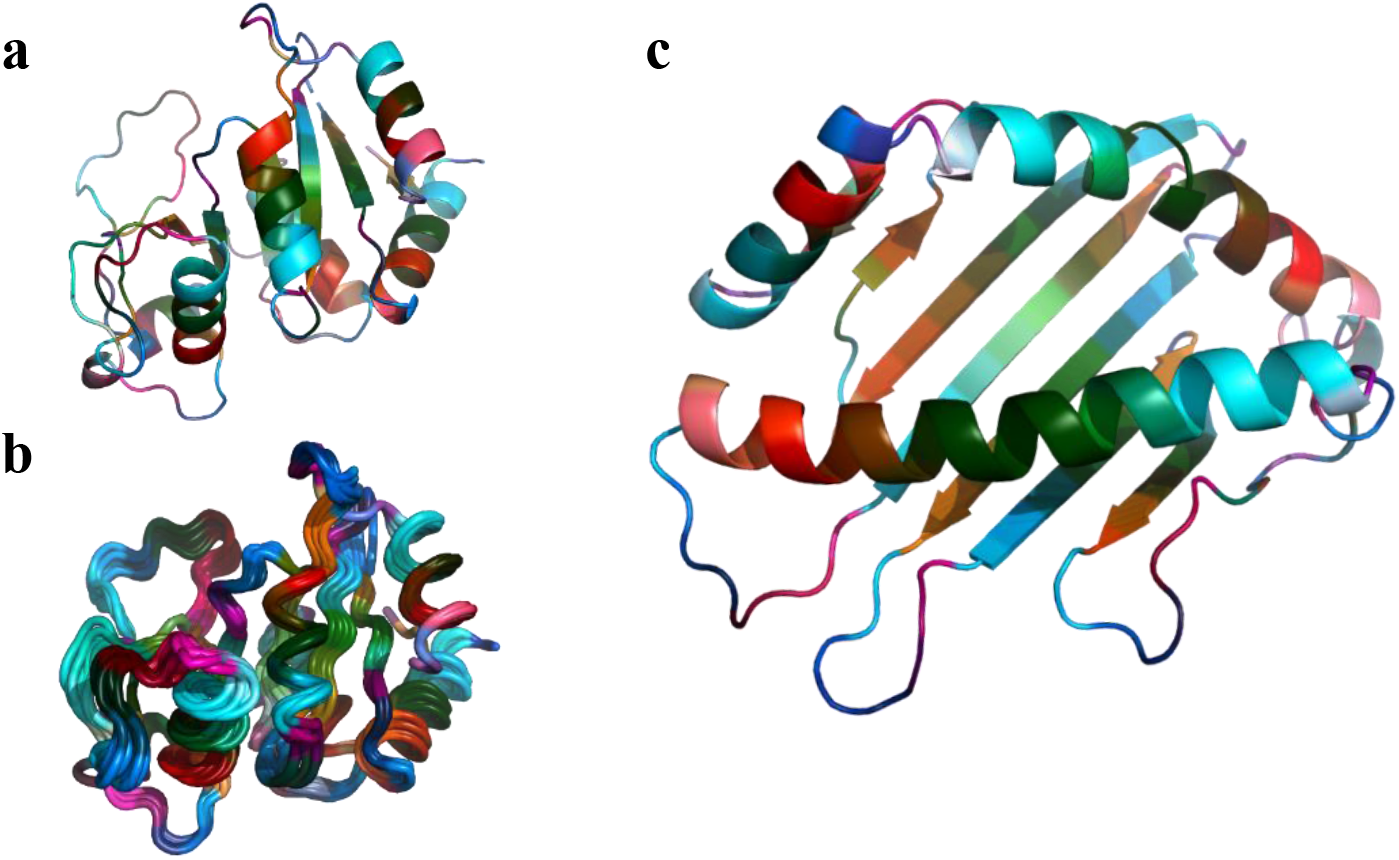
Visualization of reduced GNN_full embeddings. Dimensionality reduction is done with an autoencoder to three dimensions and translated into CIELAB colorspace. Panel (**a**) shows the embedding of 1G57. Panel (**b**) shows multiple synthologs created using this pdb as template. Panel (**c**) shows the embedding representation of MHC-I (1A1M) G-domain.

These findings demonstrate that the embeddings can differentiate between individual SSEs as they appear to begin with cyan color at the beginning of each SSE. Additionally, the embeddings encode some relative positional information over the individual SSEs, as the colors of the gradient are the same for identical SSEs, but more spread out depending on their size compared to other SSEs. For example, the long helix in MHC-I has longer stretches of cyan, green, and red compared to shorter helices but still share the same gradient. The embeddings are also able to incorporate SSE-type information, as the different types of SSE still show some differences, despite the overall’s similarities. Furthermore, the specific conformation is taken into account, as the helices of both 1G57 and 1A1M (MHC-I) have similar color embeddings but differ depending on their position in the whole protein. Notably, the embedding is robust to small rotational variations in the overall structure, as seen in the multiple variations of the same structure (Figure 8b).

Based on these changes compared to the embeddings from GNN_basic, it seems that the prominence of positional cues increased relative to the SSE type. This observation suggests that the information presented to the network influences the type of encoding that is prioritized.

### Similarity map and alignment

Since the training was conducted solely on synthetic data, we proceeded to evaluate the performance of the trained GNN on real data and compare the results with other state of the art structural alignment tools.

#### Simple case

Figure 10 showcases the alignment result between the closely related proteins 4NDK and 6M9X. While the alignment is deemed successful, it is important to note that the similarity matrix does not yield a perfect correspondence among equivalent residues. Ideally, we would expect to observe similarity primarily along the diagonal region. However, despite this limitation, there is still a significant amount of signal available to accurately superimpose the two proteins.

**Figure 10.**
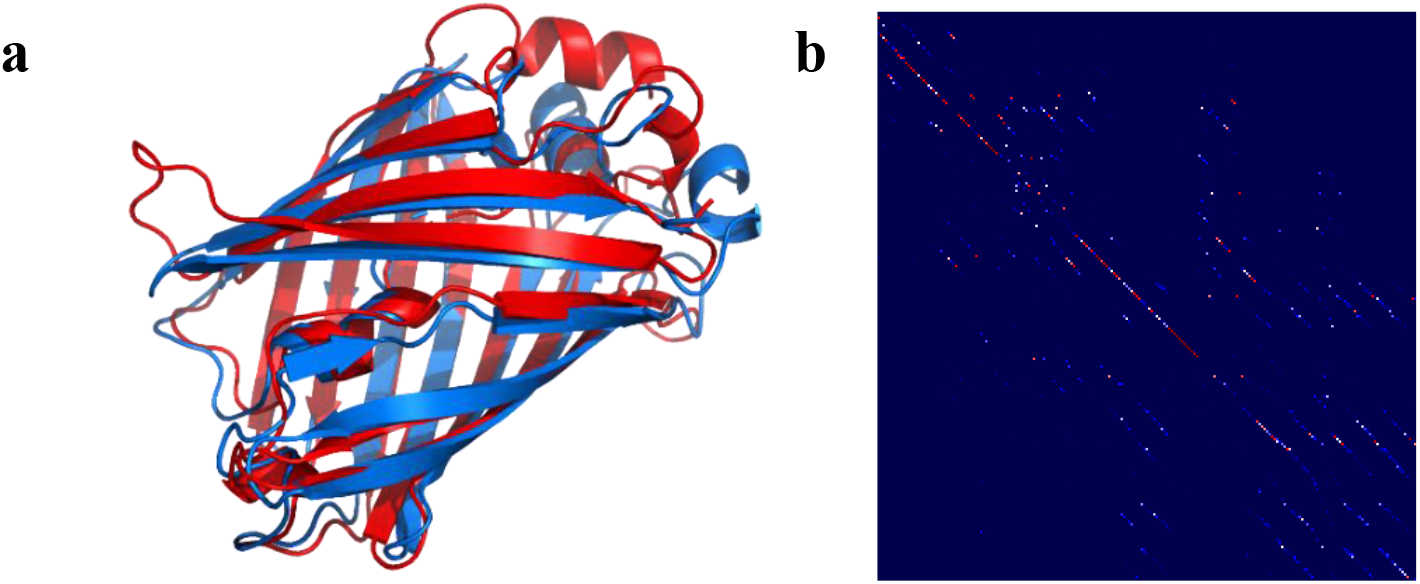
Alignment of two homologs. Homologous proteins are shown in panel (**a**) (red: 4NDK; blue: 6M9X), distance matrix between embeddings in panel(**b**). Each pixel of (**b**) represents the similarity between two residue embeddings, with red indicating high similarity and blue indicating dissimilarity The sequences are arranged starting from the top-left corner and proceeding downwards and to the right.

#### Elongation, Deletions and Insertions

While natural homologous proteins can display variations in the number and types of SSEs, our augmentation techniques focused solely on translational/rotational operations and residue substitutions. Interestingly, our structural protein alignment method successfully generalized to natural homologs that exhibited differences in SSEs, sizes, and the presence of indels, as demonstrated in the alignment of proteins 4ROF and 1HFE (Figure 11). Protein 4ROF showcases an elongated alpha helix and beta sheet, whereas protein 1HFE features a longer beta sheet located at a distinct position (indicated by purple asterisks in Figure 11). Additionally, both proteins exhibit an indel in the form of an alpha helix at another position (indicated by red asterisks in Figure 11). The ability to align homologs with diverse SSE compositions and sizes, even in the absence of such instances in the training data, underscores the robustness of the amino acid embeddings. These embeddings effectively capture local representations capable of accommodating unencountered global variations. Additional testing indicated that our method also generalized to multi-domain structures and complexes, suggesting that the learned embeddings are invariant to protein size.

**Figure 11.**
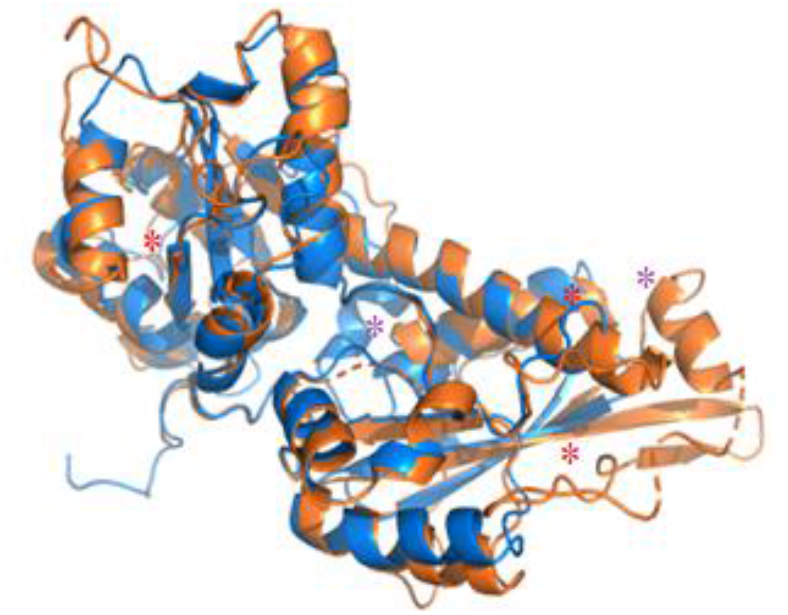
Two homologous proteins (orange: 4ROV; blue: 1HFE) with insertions of residues and SSE elongations and insertions aligned by our method. Purple asterisks indicate indels of SSEs, whereas red ones indicate elongation.

#### Comparison to state-of-the-art

The following results demonstrate the competitiveness of our GNN-based approach, where we trained the network to generate embeddings and indirectly learn structural alignment using synthetic data. As will be shown, our approach proves to be on par with, and in some cases even outperforms, other state-of-the-art structural alignment methods.

In Figure 12, we showcase an example alignment, comparing our approach to the Pymol ‘super’ method, by superimposing the G-domains of MHC-I (1A1M) and MHC-II (4I5B). These two proteins share structural similarity despite having vastly different sequences. It is noteworthy that the G-domain of MHC-II consists of two chains, while that of MHC-I has only a single chain. As depicted in Figure 12, our method exhibits significantly better overlap in both the sheets and the helices compared to the Pymol ‘super’ method despite it being specifically designed for aligning structures with low sequence similarity.

**Figure 12.**
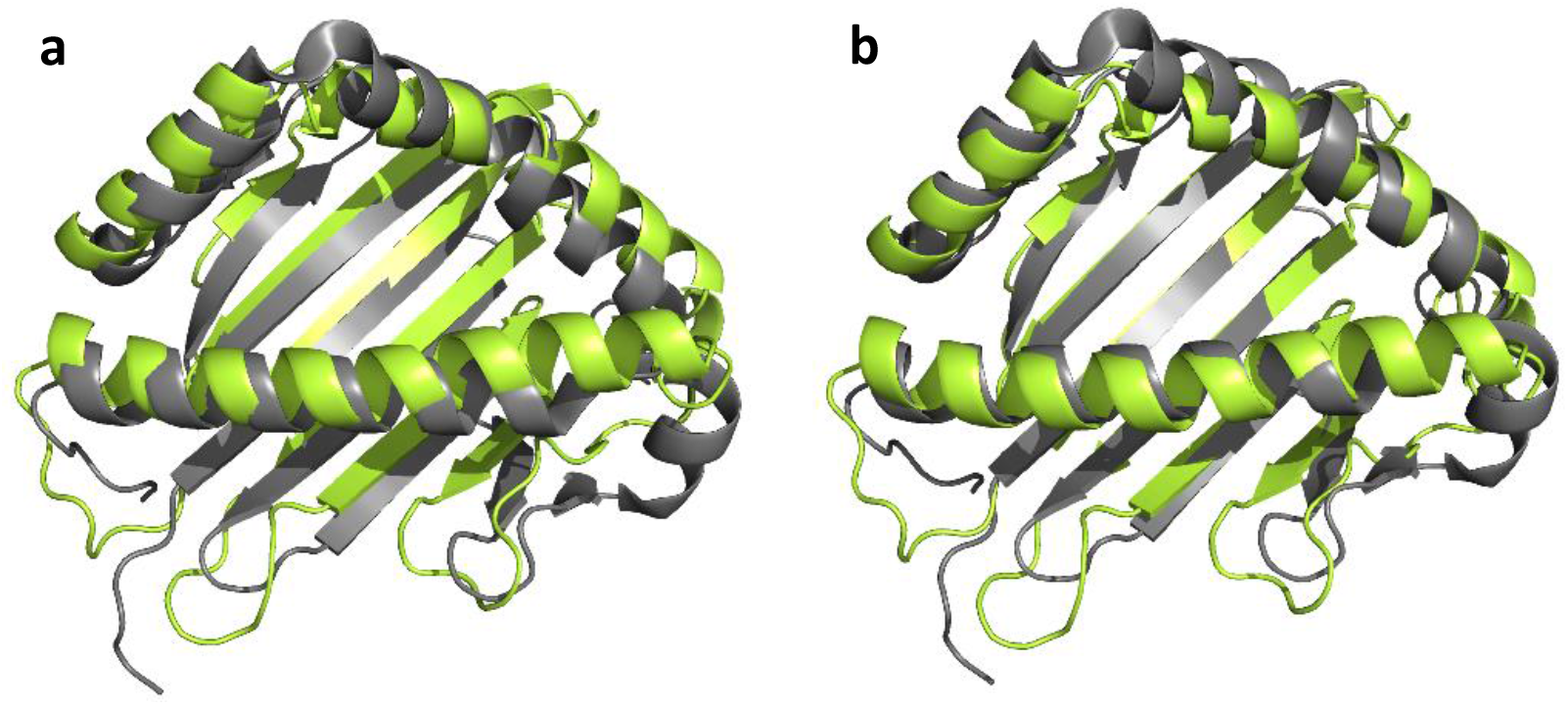
Comparing alignment between Pymol super method (**a**) and vanGOGH (**b**) on the G-domains MHC-I in green (1A1M) and MHC-II in grey (4I5B).

As aligning small proteins can pose challenges for traditional alignment algorithms, we selected proteins 1SS6 and 1VAZ (Figure 13) to compare our method with multiple top state-of-the-art methods. To evaluate the results objectively, we sought the expertise of a renowned protein expert to identify the best alignment, without disclosing the tools used for the alignments. Our method was declared to result in the best alignment.

**Figure 13.**
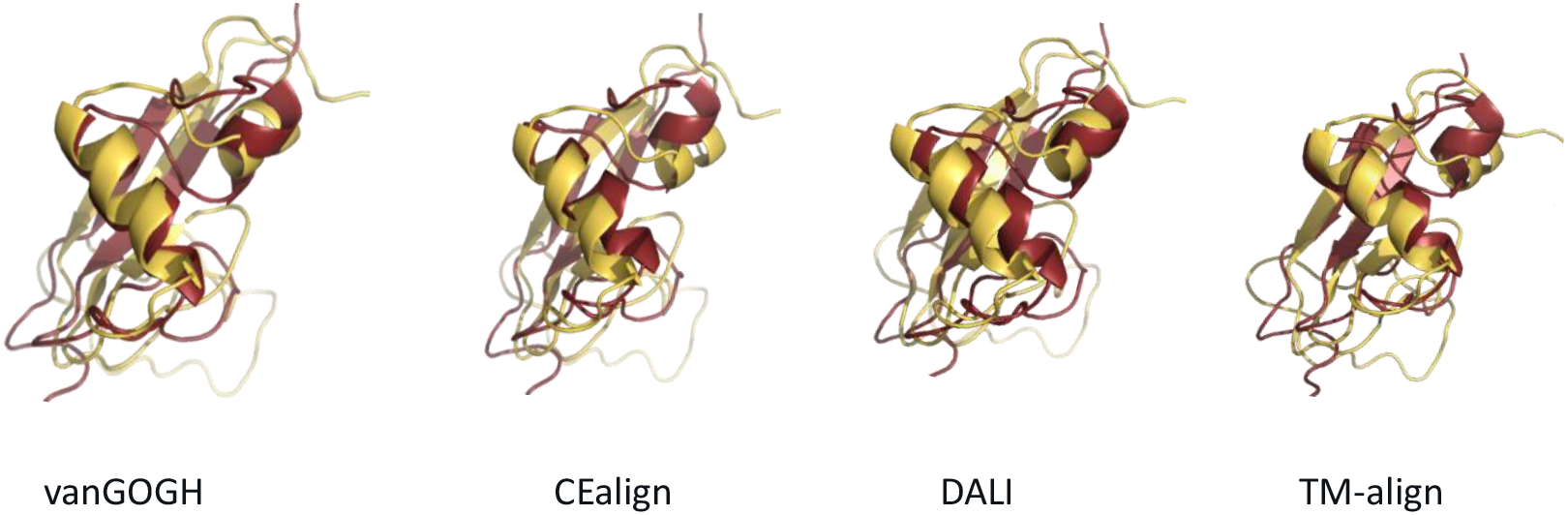
Comparison of superposition of 1SS6 (yellow) and 1VAZ (red) by our method and three state-of-the-art tools.

## Conclusion

Our study utilized a graph neural network to learn embeddings for protein structures, focusing on backbone residues for identifying corresponding residues in homologs. We trained the GNN on synthetic homologs with features designed specifically for the task, and tested it on naturally occurring homologs. The use of protein folds as training units facilitated the capture of relevant structural variations that generalized well to larger structures.

The SCOP database, while valuable, presented challenges due to its uneven distribution of domains per fold, affecting data accuracy. To counter this, we used a weighted sampling approach and introduced synthologs, creating more data for underrepresented folds and overcoming the absence of homologs in some folds.

By using synthologs, we were able to generate perfect labels for alignment training, eliminate potential biases from natural homologs, and prevent data leakage between testing and training data. This indirect method allowed us to align proteins without the need of labeled data, which further reduced potential bias. Furthermore, the probability-based alignment approach provided a natural attention mechanism for similarity scores.

The results achieved demonstrated the potential of using synthetically created homologs for protein structure alignment and the benefits of integrating domain knowledge through augmented data. Future work could include conducting a more thorough ablation study to assess the importance of amino acid features for local embeddings.

## Acknowledgments

The authors extend their appreciation to Dr. Hanka Venselaar and Prof. Dr. Peter-Bram ‘t Hoen for their valuable comments and insights. Special thanks are also extended to Prof. Dr. Gert Vriend for sharing his expertise on protein structures and conjuring the term ‘synthologs’.

